# Multiplexed, sequential secretion analysis of the same single cells reveals distinct effector response dynamics dependent on the initial basal state

**DOI:** 10.1101/511238

**Authors:** Zhuo Chen, Yao Lu, Kerou Zhang, Yang Xiao, Jun Lu, Rong Fan

## Abstract

The effector response of immune cells dictated by an array of secreted proteins is a highly dynamic process, requiring sequential measurement of all relevant proteins from single cells. Herein we show a microchip-based, 10-plexed, sequential secretion assay on the same single cells and at the scale of ~5000 single cells measured simultaneously over 4 time points. It was applied to investigating the time course of single human macrophage response to Toll-like receptor 4 (TLR4) ligand lipopolysaccharide and revealed four distinct activation modes for different proteins in single cells. In particular, we observed that secreted factors regulated by transcription factor NFkB (e.g., TNF and CCL2) predominantly show on-off mode over off-on mode. The dynamics of all proteins combined classified the cells into two major activation states, which were found to be dependent on the basal state of each cell. Single-cell RNA-Seq was performed on the same samples at the matched time points and further demonstrated at the transcriptional level the existence of two major activation states, which are enriched for translation vs inflammatory programs, respectively. These results showed a cell-intrinsic heterogeneous response in phenotypically homogeneous cell population. This work demonstrated the longitudinal tracking of protein secretion signature in thousands of single cells at multiple time points, providing dynamic information to better understand how individual immune cells react to pathogenic challenges over time and how they together constitute a population response.

## INSTRODUCTION

The advent of high throughput single-cell transcriptomic analysis has enabled global unbiased analysis of gene expression in thousands of individual cells and has revolutionized how we study the biological mechanisms at the whole organism level and the complex physiological systems such as the immune system^1 2 3^. However, accessing protein information from individual cells has been more difficult due in part to the lack of genome-wide amplification method as in nucleic acid analysis for the protein counterpart, pointing to the need to develop new technologies to collect protein information from individual cells efficiently and accurately. It is also shown in a host of studies that transcriptional and protein-level data usually show poor correlation^4^, highlighting the importance of integrating protein-information with the single-cell transcriptomic result for more comprehensive understanding of the immune system.

A group of secreted proteins including cytokines, chemokines, and cytotoxic enzymes produced by immune cells play important roles in dictating their effector response and mediating collective cellular functions^5^. Innate immune cells exert the protective defense against a variety of pathogens/viruses through not only phagocytosis but also secretion of effector proteins upon activation^6 7^. Multiplexed detection of effector proteins from single immune cells is the direct measurement of functional phenotype, providing new insights to the mechanism of innate immune responses as well as potential correlates to clinical outcome^8 9^. Significant efforts have been made to characterize single-cell secretion pattern and its correlation with cell function. For example, the quality of a CD4+ T-cell cytokine response was reported to be a crucial determinant in whether a vaccine is protective^9^, which better informs disease mechanism and drug/vaccine development^10^. Currently, most commonly used tools for multiplexed single cell secretion analysis includes ELISpot (Fluorospot)^11^, intracellular cytokine staining (ICS) flow cytometry with either fluorescence or mass spectrometry (CyTOF) detection^12 13^. Some other multiplexed proteomics profiling methods with lower throughput were also reported^14 4 15^, for example, Herr et al. developed single-cell western blotting with ~11 protein targets detected per cell after multiple fluorophore bleaching-restaining cycles ^15^. However, all these high-plex methods could not retain live cells after the assay, which make them impossible to track the same single cells at different time points but also measure a whole panel of protein secretions^16^. And such type of dynamic assay can provide unique information which inspires deeper insights resolving how immune cells respond and progress upon stimulations^17 18 19^. With the development of micro-technology, a nano-well-based microengraving assay was reported, in which sequential release of cytokines from polyfunctional human T cells can be captured^20 21^. However, the highest degree of multiplexing is four proteins, which is insufficient to dissect the full functional spectrum of heterogeneous immune cells^10 22^. Although an integrative microfluidic-based device was developed to probe single-cell multiplexed input-output dynamic, the cell number to be probed is low (only dozens) and device manipulation is highly complex.^23^ To characterize the dynamic, full-spectrum secretion information from large quantities of individual cells to understand immune function diversity, it is desirable to develop a method to measure the same single cells over multiple time points (e.g., ~4 or more) for a panel ~10 or more protein secretions and such data can be obtained from a large number (e.g. ~5000) of single cells while minimize the complexity of device handling.

Here, we described a single-cell microchip which allows for high throughput (~5000 single cells), multiplexed (~10 proteins), sequential (4-5 time points) secretion analysis of the SAME single cells. It was used to profile homogeneous human macrophages and revealed inherently heterogeneous responses over time upon activation with TLR4 ligand lipopolysaccharide. Importantly, we found the stimulated response exhibited two intrinsic states that appear to be associated with the basal function. Single-cell transcriptomic profiles collected at different time points from the samples stimulated in the same way also confirmed the presence of two cellular states with distinct gene expression profiles. Comparing single-cell protein secretion dynamics and transcriptome sequencing data allows for tracking the states of the same cells that confirmed the two states are dependent on initial cell states and differentially regulated by translational and proinflammatory programs.

## RESULTS

### Single-cell secretomic analysis microchip with higher throughput

The configuration of microchip platform used for dynamic multiplexed single cell assay was modified from previously reported devices^24 25 26 27^, which was comprised of two components: a high-density antibody barcode patterned glass substrate for surface immunoassay and a nanoliter microtrough array for single cell capture. Notably, we redesigned flow patterning microchip to combine spatial multiplexing (multi antibodies co-flow patterning) and spectral multiplexing (multi-color detection) (Figure 1A) in a much shorter microtroughs (~0.48 mm) to achieve significantly higher number of single cells to be assayed simultaneously (more than 5000 single cells data can be obtained in one microchip which is around ~5 fold increase compared to previous work). The flow patterning microchip for antibody immobilization consists of 120 repetitive barcodes, each of which contains 5 stripes in duplicate. The antibody stripes are 30 μm in width and the pitch size of a full barcode is 250 μm. Due to increased flow resistance with much longer channel length, the antibodies were flow patterned with two separated paths to solve this problem. We validated this new flow patterning strategy with both fluorescent-BSA and recombinant protein sandwich immunoassay to make sure the reproducibility of antibody barcode is adequate for single-cell experiments (**Supplementary Figures 1-3**). PDMS cell capture chip (Figure 1B) was designed according to the dimension of flow patterned antibody microarray such that it is long enough to contain at least a full set of barcodes, thereby eliminating the need for precise alignment of the antibody barcode slide and the microtrough array PDMS slab. With this design, more than 5000 single cells data (around 30% of total number of microtroughs (n=18000), Figure 1C) can be detected in one microchip with minimal sacrifice of parameters to be plexed. For example, up to 15 different proteins can be profiled at the same time if three color detection strategy were employed.

**Figure 1.**
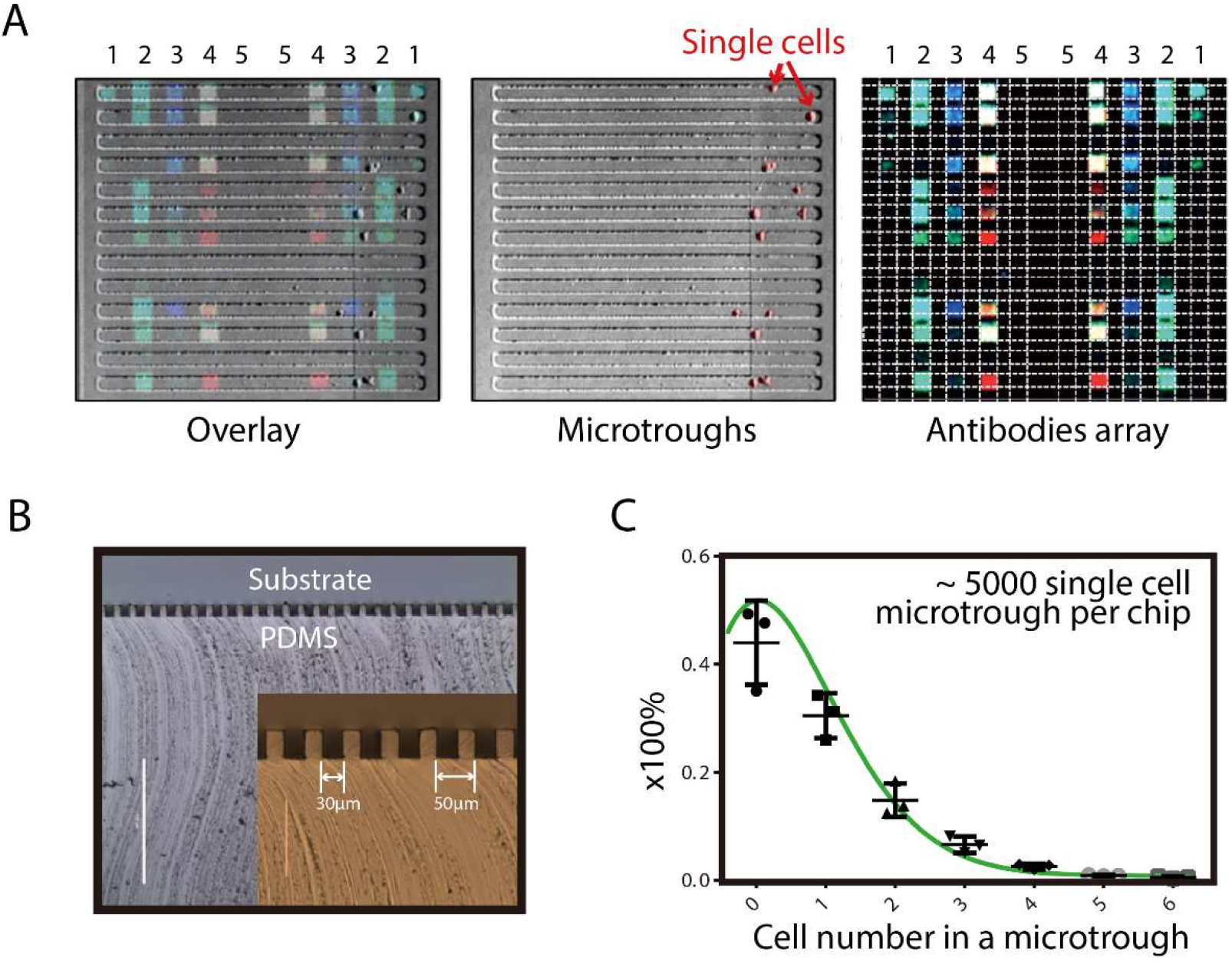
Single-cell secretomic analysis microchip with higher throughput. a) Images showing the captured single cells, corresponding three-color fluorescence detection results (Red: 635 nm, green: 532 nm and blue: 488 nm) and their overlay. b) Cross-sectional view of microtroughs. Inset: enlarged view of the microtrough cross-sections. The width of each microtrough is 30 μm. c) Distribution of the number of cells per microtrough under optimized cell loading conditions (cell density: 0.2 million/mL, loading volume: 200 μL, loading time: 5 min), which reveals around 30% of microtroughs would be occupied by single cells.

### Multiplexed, sequential secretion analysis from the same single macrophages reveals heterogeneous cytokine secretion dynamics

One unique feature of our single-cell assay platform is that cells assayed are alive and still isolated in defined locations (specifically for adherent cells). High reproducibility in protein secretion frequency is also validated (**Supplementary Figure 4**). All these make it possible to accurately and dynamically track the secreted proteins from the same single cells at different time points. Briefly, after measuring protein secretion from single cells over a period of time, we removed the antibody barcode slide that captured the basal secretion profile, and then replaced with a new antibody barcode slide to measure protein secretion from the same single cells for another period of time, during which stimulation reagents can be added, withdrawn, or combined, permitting flexible design of the experiment to perturb cell signaling but keep track of the same single cells over time. Repeating this process will lead to the measurement of single-cell protein secretion dynamics (Figure 2A). The PDMS microchip for cell capture is oxygen plasma treated for 1 min just before single cell experiment to make its surface hydrophilic to enhance cell adhesion and minimize nonspecific protein adsorption^28^. When changing a new antibody array slide, the PDMS microtrough chip was rinsed three times (including washing and incubation for 2 minutes in each step) with fresh medium to wash out residual secreted proteins. This also removes detached cells that may dislodge to neighboring microtroughs. Figure 2B shows that 56% of macrophage cells could be retained after the removal and replacing with a new antibody slide. Figure 2C shows a representative single cell that was retained in the same microchannel throughout the entire secretion dynamics experiment and the corresponding secretion patterns.

**Figure 2.**
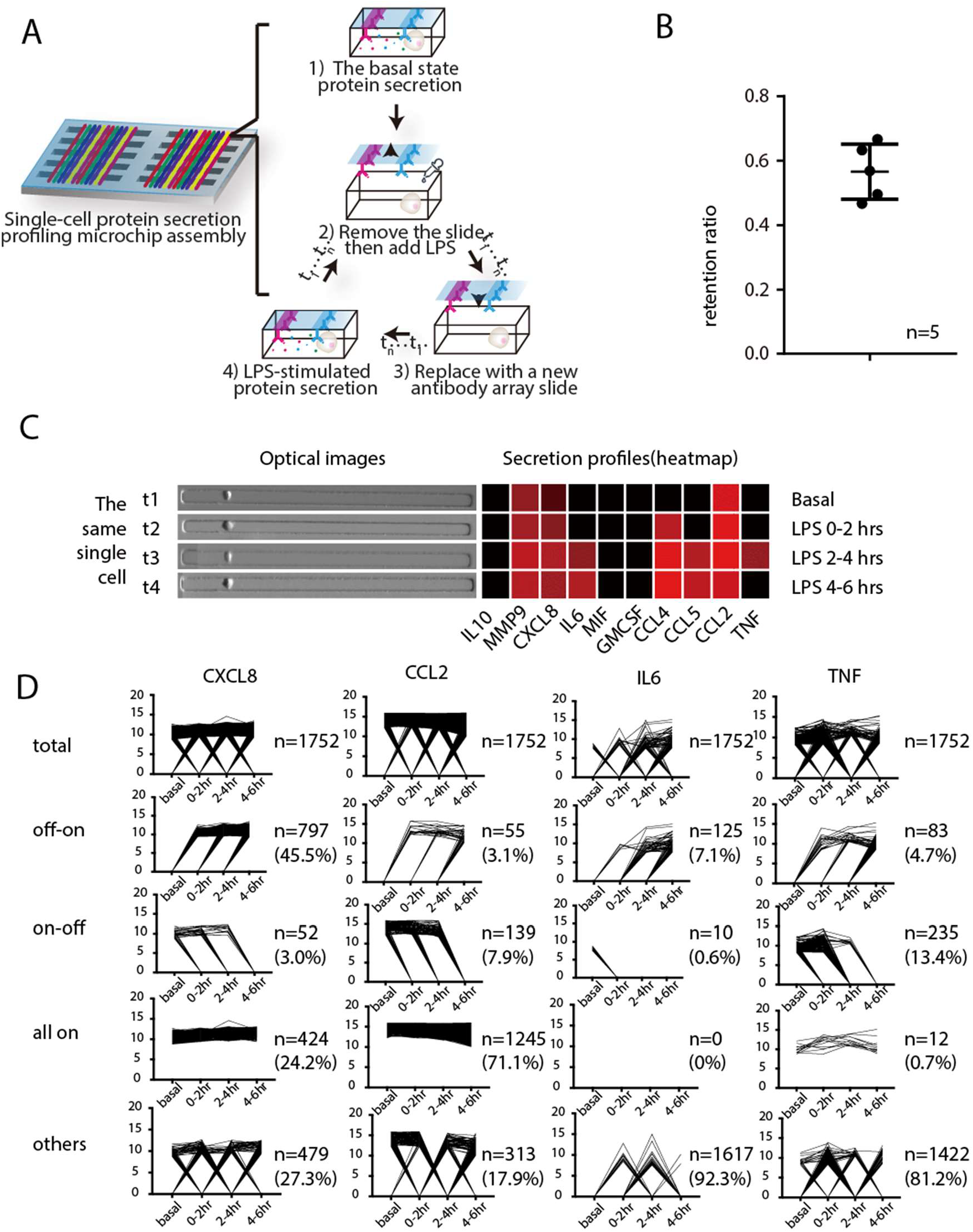
High-throughput, multiplexed, sequential secretion analysis from the same single cells. A) Schematic illustration of the procedure for tracking the secretion from the same single cells at different time points. Single-cell secretions were firstly profiled at the basal state, then LPS was added to induce macrophage responses to TLR-4 activation. The time course of LPS-induced activation is measured sequentially within different time windows; B) Characterization of single cell retention efficiency with human macrophage cells after a new antibody glass slide was changed to PDMS microchamer array (n=5); C) Representative single cell and its secretion pattern of 10 proteins from the same single cell at 4 time points, before and after LPS treatment; D) Line graphs showing the dynamic change of different secreted proteins (CXCL8, IL-6, CCL2, TNF) from1752 single cells, which were classified into four patterns based on their protein secretion dynamics: on-off, off-on, all on or others.

We applied this platform to investigate the dynamics of U937 derived macrophages in response to Toll-like receptor 4 (TLR4) ligand lipopolysaccharide, which simulates the innate immune response to Gram-negative bacteria^29 30^. The U937 monocyte was differentiated into macrophages by 50 ng/mL PMA for 48 hours and confirmed by CD14 surface marker staining (**Supplementary Figure 5**)^31 32^. The antibody pairs used in this study (**Supplementary Table 1**) were validated with corresponding recombinant proteins for crosstalk reactivity to ensure technical validity. We also obtained the titration curves, which demonstrated the feasibility of comparing the amount (or concentration) of secreted proteins semi-quantitatively (**Supplementary Figure 6&7**). After assaying for 4 time points, a set of data comprising 1752 single cells, each of which has data for a full time course and simultaneous detection of 10 secreted proteins was successfully obtained. It allowed to compare the dynamic change of different secreted proteins of each cell at basal state and with LPS stimulation at different time points (Figure 2D). For example, TNF was firstly secreted upon LPS stimulation and the intensity decreased after a 2hr period. Other effector proteins like IL-6 and IL-10 were secreted afterwards, which is in agreement with previous reports^33 34^.

The dynamic change of each protein in every single cell during the time course can thus be visualized in Figure 2D by connecting their respective detection results at each time point. All single cells (n=1752) were classified into four subgroups according to their secretion dynamics: on-off, off-on, all on and others. The secretion dynamics designated as “on-off’ means the protein secretion is active at early time window (s) but stopped in a later period; “off-on” means the secretion was not detectable initially but became active later on; “all-on” means this specific protein was secreted continuously throughout the time course of observation; “other cases” include no secretion or oscillatory secretion patterns. It is noted that the grouping result differs with respect to specific proteins of interest. We noticed that CXCL8 and CCL2 secretions in most cells is “on” during (at least partially) the period of observation, but their dynamics are quite different: the majority of the cells are not secreting CXCL8 during basal state and start turning on CXCL8 secretion after being stimulated. However, the timing for them to turn on CXCL8 secretion is heterogeneous, with comparative numbers of cells turn on CXCL8 secretion between 0-2hrs, 2-4hrs, and after 4hrs after being stimulated. Quite differently, CCL2 secretion in the vast majority of cells (71.1%) are kept in the “on” mode independent of being stimulated or not, with a small fraction (7.9%) of cells turn CCL2 secretions off throughout the period of observation. We then specifically looked at two factors (IL6 and TNF) regulated by the transcription factor NFkB, which is one of the most critical and widely recognized transcription factors regulating inflammatory responses in immune cells. According to our data, a majority of the cells are classified as “others”, meaning not secreting or secreting in oscillatory manner. This is consistent with the previous reports of NFkB oscillation during transcriptional regulation. However, we also observe a small portion of cells with stable “on-off’ or “off-on” secretion patterns, breaking the rule of oscillation. Interestingly, within the population with stable secretion patterns, there are far more cells showing “off-on” pattern versus “on-off’ pattern in IL6 secretion (7.1% vs. 0.6%) while the trend is reversed in the case of TNF secretion (4.7% vs 13.4%), suggesting cellular basis of a more transient nature of TNF response. We selected 7 proteins being secreted by noticeable number of cells and with apparent secretion alterations during the course of measurement for this analysis. Figure 2D shows the single-cell secretion patterns of 4 proteins and 3 others are shown in **Supplementary Figure 8**. Previous study has revealed the stochastic and variable cell switching dynamics of NFkB in response to TNF activation at single cell level^19^. Herein we show that the transcriptional & functional output of NFkB activation such as cytokine secretion could exhibit oscillation patterns but the cytokine function outputs are more diverse, which has never been observed previously.

### Responses of macrophages are intrinsically heterogeneous and dependent on the basal functional state

By combining all the single cell data at four different time points (40 proteins parameters) into a unique dynamic, multiplexed single cell data metric, which cannot be obtained using other methods, we performed unsupervised hierarchical clustering and resolved two distinct major clusters with 1133 and 619 cells, respectively (Figure 3A), indicating the presence of a dynamic heterogeneity within a phenotypically homogeneous macrophage population in response to LPS. Cluster 1 (red label) cells are more active in secretion than cluster 2 (black label) cells not only after stimulation but also in the basal state. We applied a high-dimensional data analysis tool viSNE to visualize multiplexed single-cell data^35^, from which two similar clusters can be obtained too with 943 and 809 cells, respectively (Figure 3B). Further analysis identified a ~70% overlap in both clusters using those two clustering methods, indicating the robustness of resultss regardless of clustering algorithms. The expression of specific proteins in single cells within each cluster (e.g., CCL4, TNF in Figure 3C, and **Supplementary Figure 9**) at different time points can be shown in viSNE plots, from which we confirmed the cells more active at basal state are more prone to secreting more proteins upon pathogen stimulation. This phenomenon was not observed previously, highlighting the value of high throughput dynamic detection platform with single cell resolution in resolving how biological systems respond and evolve.

**Figure 3.**
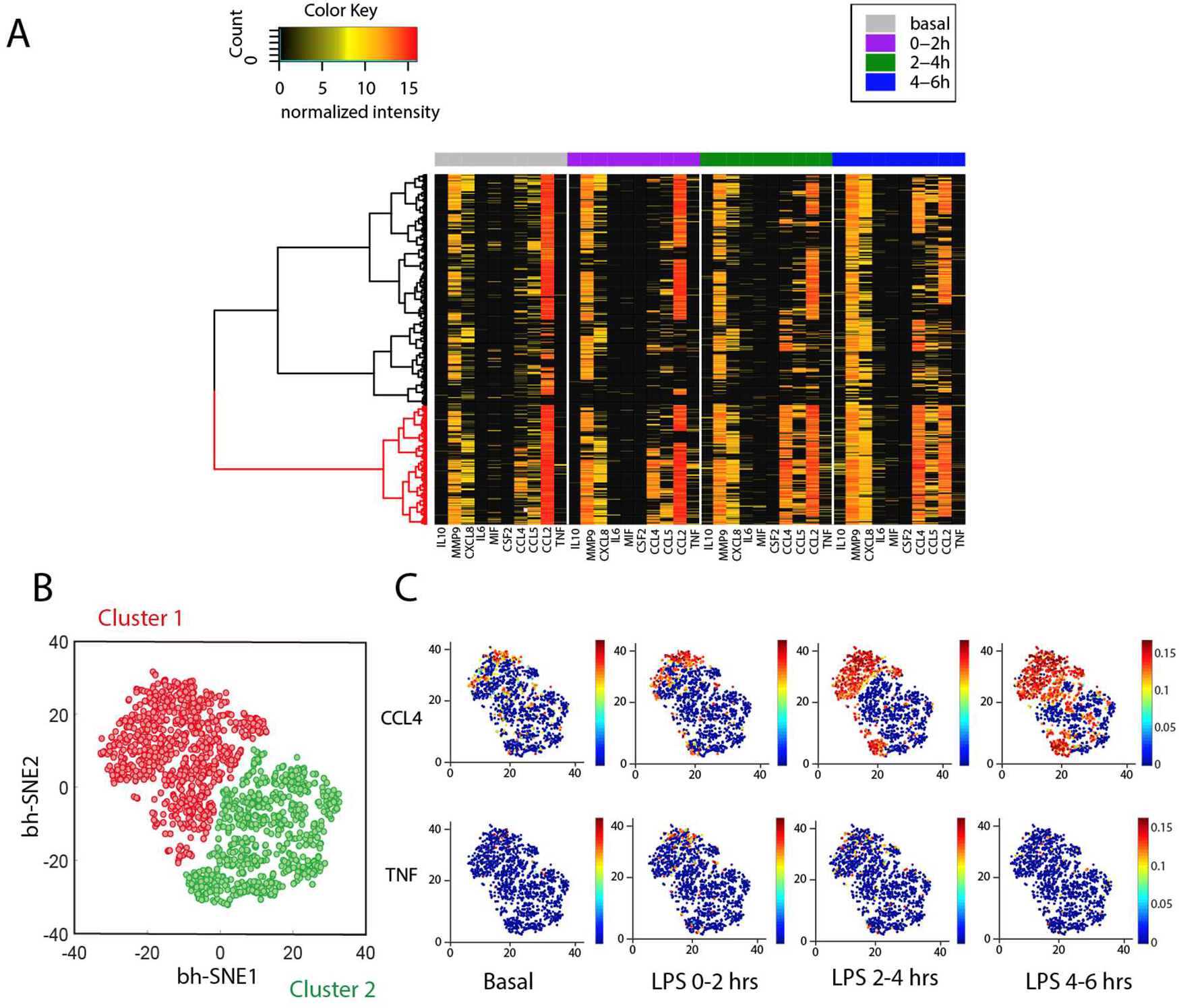
Dynamic heterogeneity of U937 derived macrophages in response to TLR 4 ligand LPS. A) Heatmap resolving two clusters exhibiting different activation dynamics in response to LPS stimulation. Each row represents a complete protein profile from a single cell and each column is a protein of interest; B) viSNE analysis reveals two clusters based on their dynamic functional proteins profiles. C) Distribution of individual proteins (CCL4, TNF as examples) in viSNE maps colored by signal intensities in two clusters at different time points. (viSNE plots for distribution of all proteins can be found in **Supplementary Figure 7**).

Quantitative analysis of single-cell polyfunctionality (the ability of a cell to co-secrete multiple cytokines and chemokines simultaneously^36^) was performed and compared between the basal state and the stimulated states, which showed an increase of highly polyfunctional cells upon stimulation (Figure 4A). It is previously reported that polyfunctionality index (PI) serve as an effective parameter to numerically evaluate the degree of polyfunctionality from multi-dimensional single cell data, permitting more sophisticated statistical analysis^37^. We applied a previously described approach (detailed in supplementary methods) to calculate PI of single cells^37^. Consistent with previous result, we see an increase of PI immediately after stimulation comparing with cells in resting state (Figure 4B). Interestingly, PI remains constant in the early stage of activation but increased after 4 hours being stimulated. This suggests that macrophages increase polyfunctionality immediately (<2hr) after being stimulated but it takes several hours (>4hr) for them to further increase polyfunctionality an enter a more advanced activation state. We then compared polyfunctionality of the SAME individual cells between different time points and looked for correlative patterns (Figures 4C, 4D &4E). Interestingly, the resulting averaged polyfunctionality of stimulated single cells showed a linear correlation with the basal state polyfunctionality (R^2^=0.99, 0.96, 0.98 at 0-2 hrs, 2-4 hrs, 4-6 hrs respectively after LPS stimulation). This result means that the higher the numbers (types) of proteins that a cell is secreting in its basal state, the higher numbers (types) of proteins it will likely secrete after being stimulated, and the basal state secretion activity can be taken as an indication of its later activity, suggesting the activation potential of macrophages likely pre-determined even before the arrival of external stimulation and is encoded in the basal state intrinsically (Figures 4D & 4E).

**Figure 4.**
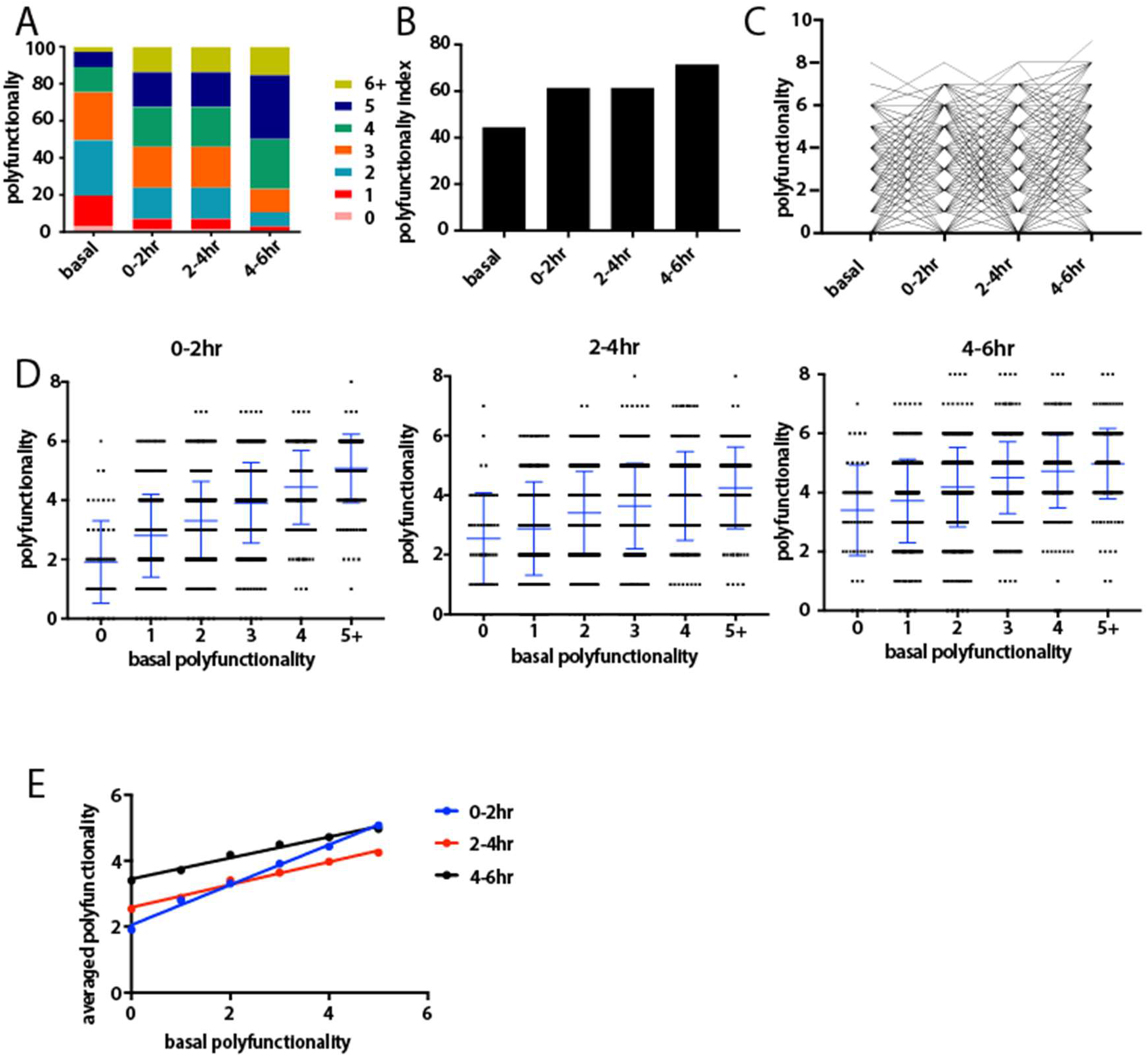
Polyfunctionality analysis of human macrophages. A) Distribution of polyfunctionality among macrophages of each time window (basal state included). B) Gradual increase of polyfunctionality index (PI) along the immune activation process. (C) Tracking polyfunctionality of each individual macrophage over time. D) Distribution of polyfunctionality of activated macrophages of each condition compared with basal polyfunctionality. The numbers of single cells with different poly-functionality (0, 1, 2, 3, 4, >5) are 94, 360, 548, 427, 228 and 95 respectively. The p value by t test between neighbors were <0.05 if not stated otherwise (ns: no significant). E) Linear correlation between later averaged polyfunctionality and the basal polyfunctionality.

### Single-cell RNA-Seq confirms two major clusters with distinct gene expression profiles

We also performed single-cell RNA sequencing analysis on the same samples (both basal and stimulated for different times) in order to compare to single-cell protein secretion data and further investigate the mechanisms underlying the observed heterogeneous states. A massively parallel 3’ mRNA capture and barcoding in droplets was used and the library construction was similar as previously described^38^. The raw sequencing data were processed and analyzed for quality assessment (**Supplementary Figure 10**) and the generation of single-cell gene expression matrix. Single-cell transcriptome data of all samples including both basal and activated macrophages of each time point was combined and analyzed with the R package Seurat. Graph-based clustering (resolution=0.15) identified 3 clusters when all the samples combined, named clusters 0, 1 and 2 (Figures 5A & 5B). Cluster2 largely overlaps with the basal or resting macrophages before activation, while clusters 0 and 1 largely correspond to activated macrophages, indicating the existence of two major distinct activation states in agreement with single-cell protein secretion data. Those two activation states consistently exist in samples of all time points post activation, marking a highly consistent and possibly stable dichotomy in activation states of human macrophages in response to LPS and the top ranked signature genes defining each cluster were identified (Figure 5C & **Supplementary Figure 11**). Although mRNA data of activated macrophages supports protein data well, partially confirms our argument that cellular states are possibly pre-determined and stable, we noticed that the separation of resting macrophages clearly into two clusters is not seen in the transcriptomic profile. In figures 5A, the observation of three clusters supports the two-state activation model derived from single-cell protein sequential secretion. As expected for whole transcriptome analysis, tSNE separates basal from stimulated cells. The latter however exhibits two clusters, suggesting two major activation states as revealed by single-cell protein data. Figure 5B further confirmed the gradual change of cell states over time and two major states are always present at any given time points. Interestingly, the basal state cells (cluster 2) already showed “bifurcation” toward two different activation states as defined by Clusters 0 and 1. All these are in agreement with single-cell protein secretion data and support the conclusion that the activation shows two distinct states.

**Figure 5.**
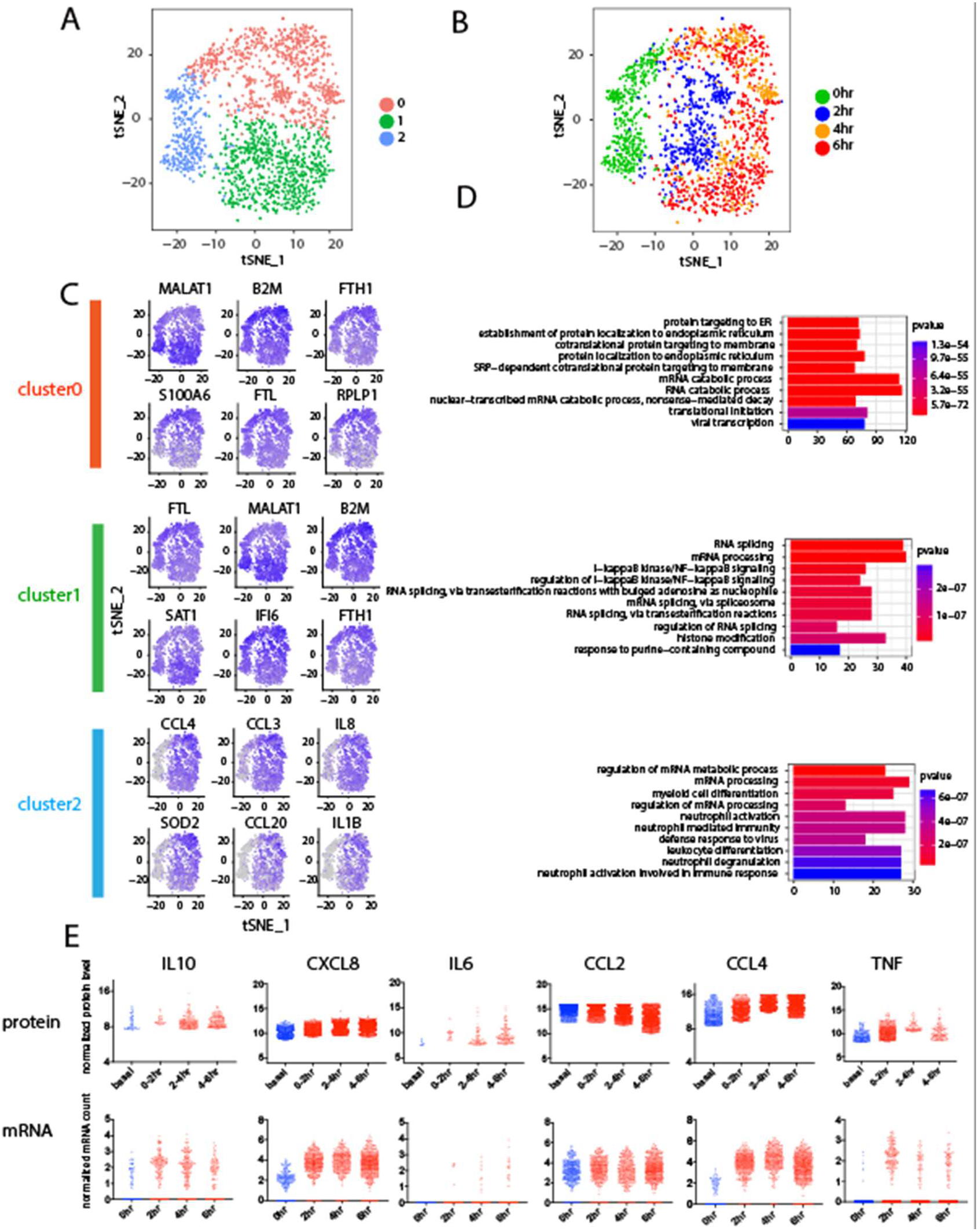
Single-cell RNA Seq of U937-derived macrophages during dynamic TLR4 immune activation. A) and B) tSNE plots showing dynamic formation of two cellular sub-states in activated macrophages. Cluster2 overlaps with resting macrophages while clusters 0 and 1 are distinct sub-states of activated macrophages. C) tSNE feature plots showing expression of marker genes of each cellular sub-state. D) Gene Ontology (GO) term enrichment analysis reveals differential expression of immune activation and ribosomal genes in two sub-states among activated human macrophages. E) Comparative analysis of single-cell mRNA and protein secretion data.

We then asked what gene pathways and biological processes underlie the previously identified two activation states. Gene Ontology (GO) analysis was performed with the R package GAGE using the gene lists and P values generated from the Seurat program as described above, and the marker genes of each of the 3 clusters were used. Surprisingly, the GO terms highly enriched in cluster2 involves mRNA processing, which indicates that mRNA processing related genes are highly expressed in the macrophages to prepare them for activation. Furthermore, we found that the GO terms enriched in the two clusters of activated macrophages (clusters 0 and 1) involve protein translation (ribosomal genes) and inflammatory response (NFkB pathway, etc.), respectively (Figure 5D). The anticorrelation relationship between the activity of protein translation and the ability to mount inflammation appears to be mutually exclusive in a single macrophage cell such that each cell has to opt for only one specific state. However, it is unclear if this is a stochastic process or pre-determined by the initial state according to single-cell transcriptome data. The protein secretion dynamics measured on the same single cells suggests the activation states are dependent on the basal state of each macrophage.

We further conducted integrative analysis of single-cell transcriptome and protein secretion data side-by-side, and observed gene-dependent correlations between single-cell RNA and protein profiles. Figure 5E shows 6 immune function genes measured at both transcriptional and protein levels and other genes of interest are shown in **Supplementary Figure 12**. The integrated single-cell transcript/protein data can be generally classified into three groups. First, highly correlated genes, including IL6, CCL2 and TNF. Second, genes with consistent dynamics but distinct expression levels, including CXCL8 and CCL4. Third, genes with consistent expression levels but distinct dynamics. IL10 is an example in this category (Figure 5E). IL10 mRNA peaks at 2 hours, however, IL10 protein secretion reaches maximum between 4-6 hours. This can be explained by the time lag between transcription and translation. Although the mechanism requires further investigation, our results shows consistency and discrepancy in a gene-specific manner between single-cell mRNA and protein data, which is in agreement with previous reports^4^, highlighting the value to integrate information from multiple omic levels to yield a comprehensive biological picture.

## DISCUSSION

We reported a highly multiplexed (*≥* 10) single-cell sequential secretomic profiling platform to evaluate the dynamic immune response from the same single cells at different time points. With this platform, we found human macrophages in response to TLR4 stimulation exhibit two intrinsically distinct states, which were correlated with the basal functional state in each single cell. We also applied single-cell RNA-Seq analysis to confirm the existence of two major states at the transcriptional level and further revealed that the two activation states are differentially dictated by the expression of protein translation genes and inflammation-related genes, respectively. It has not been possible to obtain this type of comprehensive protein secretion dynamics information of individual cells using any other methods. An even higher degree of multiplexing can be obtained if additional fluorescence channels were used. While this work focused on human macrophages, this approach can be readily applied to other adherent cell types such as epithelial cells, dendritic cells, fibroblasts, etc, making it a versatile platform well suited for much broader applications^39 40 41^. This platform will provide an accessible tool for more in-depth and comprehensive monitoring of cell function and the cognate proteins. It may have the potential to further evolve and be adopted in clinical and pharmaceutical studies to evaluate cellular function heterogeneity or drug responses^39 42 43 44^, for example, in the immune system or tumor microenvironment.

## MATERIALS AND METHODS

### Fabrication of antibody barcode array chips and microtrough array chips

The mold for both the flow patterning PDMS replica and the subnanoliter microarray are silicon master etched with deep reactive-ion etching (DRIE). Detailed protocol has been described elsewhere^25^. The PDMS used was RTV615 from Momentive. A and B were in 10:1 ratio.

### Cell culture and stimulation

Human U937 cell line was purchased from American Type Culture Collection (ATCC) and cultured in RPMI medium 1640 (Gibco; Invitrogen) supplemented with 10% FBS (ATCC). The U937 cells were differentiated with 50ng/mL phorbol 12-myristate 13-acetate (PMA) (Fisher) for 48h, followed by culture in fresh standard medium for 24h. The cells were harvested with trypsin for single-cell experiments. Cell were challenged with 100ng/mL LPS (Sigma).

### Single-cell longitudinal multiplexed secretion assay

PDMS microtrough array was first blocked with 3% BSA solution (Sigma) for 2h and then rinsed with fresh cell medium. Cells were suspended in fresh medium at concentration of 0.2 million cells per mL, followed by the addition of LPS as described previously. The PDMS microtrough array was placed facing upward, and cell culture media was removed until only a thin layer remained on surface. Cell suspension was pipetted (200 μL) onto the microtrough array. 5 minutes later, the glass slide with antibody barcode was put on top of the PDMS microtrough array with the antibody-patterned side facing the cell-capturing chambers. Then the two parts were clamped tightly with screws using a custom polycarbonate clamping system. Number and locations of cells were confirmed by optical imaging using Nikon Eclipse Ti Microscope with an automatic microscope stage. Bright field images were obtained. The assembled microchip was placed in a standard 5% CO_2_ incubator at 37° C during the period of cell secretion. After every 2h, the microchip assembly was dissembled in fresh media and the antibody barcode slide was removed and rinsed with excessive fresh media followed by 2 minutes of incubation. After repeating this step for 3 times, the cell capture microarray chip was rinsed with fresh media with stimulants added and a new glass slide with antibody microarray is applied on the cell capture microarray to generate a new microchip assemble, followed by imaging and incubation, as described. On the other hand, the glass slide dissembled previously was developed for 1h at room temperature by introducing a mixture of biotinylated detection antibodies. The detection antibody mixture consists of the detection antibodies (SI Appendix, Table S1) at 0.25 mg/mL each in 1:200 suspension in 3% BSA. Following this step, the slide was rinsed with 3% BSA solution. The 200μL of 1:100 suspension APC dye-labeled streptavidin (Biolegend, 5μg/mL) were added onto glass slide to detect the 635-nm detection antibody group, followed by incubation of 30 min. Following the BSA blocking, the antibody barcode array slide was rinsed with 1xPBS, 0.5xPBS, and DI water sequentially, dried with forced N2 gas, and then scanned with a four-laser microarray scanner (Molecular Devices; Genepix 4200A) for protein signal detection. Microtrough array images with cell counts were subsequently matched to their protein signals for further data analysis.

### Titration experiment using recombinant proteins in microfluidic chips

The titration curves were obtained using recombinant proteins and measured on antibody barcode chips, similar as the ones used for single-cell protein secretion assay (see above). The antibody barcode glass slide was thermally bonded to a PDMS microchannel array slab and then blocked with 3% BSA/PBS solution for 1 hr. Recombinant proteins with different concentrations were introduced into different microchannels followed by incubation at 37° C for 1hr. After that, a cocktail of detection antibodies was added to complete the immune-sandwich assay using the procedure provided by antibody vendors.

### Fluorescence imaging and analysis

Genepix 4200A scanners (Molecular Devices) were used to obtain scanned fluorescent images. Two channels, 488 (blue) and 635 (red), were used to collect fluorescence signals. The image was analyzed with Genepix Pro software (Molecular Devices) by loading and aligning the microtrough array template followed by extraction of fluorescence intensity values per antibody per microtrough. Fluorescence results were extracted with the image analysis tools in Genepix Pro, and then matched to each of the microtrough array for cell counts as previously extracted from the optical images.

### Analysis of single-cell protein secretion data

Cell counting was automatically performed by a C++/QT QML software (Isospeak; Isoplexis). Protein signal data were extracted from the multicolor fluorescent images using GenePix Pro 6.1 (Molecular Devices) by aligning a microtrough array template with feature blocks per antibody per microtrough to the protein signal features. Data were extracted using the image analysis tool to gain the mean photon counts per protein signal bar per microtrough and match to the cell counts from the microtrough array. The cell counting and protein signal data were then matched based on their spatial locations. Only the 1-cell wells and their protein signals were used for downstream data analysis. 0-cell wells and their protein signals were used as on-chip controls to provide a measure of local antibody-specific background and were averaged across region on chip. We define secretion threshold of a specific factor as mean of the zero-cell wells of its corresponding antibody plus 3 times standard deviation. Values higher than the threshold are taken as “secretion” while values below it are taken as “no secretion” and are changed to 0. The thresholded data were log2 transformed using log2(x+1) before data visualization. Graphpad Prism 7 was used to generate scatter plots and line graphs. Hierarchical clustering and heamaps were performed in R. Polyfunctionality index is calculated using the following equation:

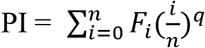

where n=6 is the cutoff in the number of functions studied, F_i_ is the frequency (%) of cells performing i functions, and q is a positive number used to modulate the differential weight assignment of each F_i_. Here q is set to 1 assuming equal weight of each F_i_.

### Single-cell RNA Sequencing

Single-cell RNA Sequencing was performed following the standard DropSeq protocol as described before in detail^45^. Microfluidic device was built following the exact original design file. Beads used in the experiment with oligo synthesize on surface was purchased from ChemGenes. Reagents used were listed in SI Appendix, Table S2. All oligonucleotides used were identical with which described in previously^45^. Sequencing was carried out using Hiseq2000 with 4 samples pooled into one sequencing lane.

### Analysis of single-cell transcriptomic data

Original fastq data was trimmed and transformed into digital expression matrix following the DropSeq data analysis pipeline described in detail elsewhere. Then a filter was applied to get rid of cells expressing fewer than 500 genes, which are likely low-quality cells. Then we applied R package “Seurat” in performing downstream statistical analysis and data visualization using default settings^46^. Marker gene lists and p values output from “Seurat” was taken as input for gene ontology and pathway analysis, using packages “Gage” and “Pathview” with default settings^47 48^.

## ACKNOWLEDGMENT

This work was supported in part by U54CA193461 (to R.F.), U54CA209992 (Sub-Project ID: 7297 to R.F.), R21CA177393 (to R.F.), National Science Foundation CAREER Award CBET-1351443 (to R.F.), NIH grants R01CA149109 (to J.L.), R01GM116855 (to Y.D. and J.L.), Connecticut RMRF grant 15-RMB-YALE-06 (to J.L.), and Sackler Institute Research Grant. Y. L. acknowledges the funding support from National Natural Science Foundation of China (Grant No. 21874133, 21605143), Youth Innovation Promotion Association CAS (Grant No. 2018217), and Dalian Institute of Chemical Physics (Grant No. SZ201601). Services provided by the NIDDK-supported Yale Cooperative Center of Excellence in Hematology assisted this study. We acknowledge the Becton Nanofabrication Center for supporting microchip fabrication and Yale Center for Genomic Analysis (YCGA) for next generation sequencing service.

## References

1. Shapiro E, Biezuner T, Linnarsson S. Single-cell sequencing-based technologies will revolutionize whole-organism science. Nat Rev Genet. 2013; 14(9):618–630. doi:10.1038/nrg3542.

2. Satija R, Shalek AK. Heterogeneity in immune responses: From populations to single cells. Trends Immunol. 2014; 35(5):219–229. doi: 10.1016/j.it.2014.03.004.

3. Papalexi E, Satija R. Single-cell RNA sequencing to explore immune cell heterogeneity. Nat Rev Immunol. 2018; 18(1):35–45. doi:10.1038/nri.2017.76.

4. Darmanis S, Gallant CJ, Marinescu VD, et al. Simultaneous Multiplexed Measurement of RNA and Proteins in Single Cells. Cell Rep. 2016; 14(2):380–389. doi:10.1016/j.celrep.2015.12.021.

5. Zhao JL, Ma C, O’Connell RM, et al. Conversion of danger signals into cytokine signals by hematopoietic stem and progenitor cells for regulation of stress-induced hematopoiesis. Cell Stem Cell. 2014; 14(4):445–459. doi:10.1016/j.stem.2014.01.007.

6. Stow JL, Ching Low P, Offenhäuser C, Sangermani D. Cytokine secretion in macrophages and other cells: Pathways and mediators. Immunobiology. 2009; 214(7):601–612. doi:10.1016/j.imbio.2008.11.005.

7. Wynn TA, Chawla A, Pollard JW. Macrophage biology in development, homeostasis and disease. Nature. 2013; 496(7446):445–455. doi:10.1038/nature12034.

8. Almeida JR, Price DA, Papagno L, et al. Superior control of HIV-1 replication by CD8 + T cells is reflected by their avidity, polyfunctionality, and clonal turnover. J Exp Med. 2007; 204(10):2473–2485. doi:10.1084/jem.20070784.

9. Darrah PA, Patel DT, De Luca PM, et al. Multifunctional TH1 cells define a correlate of vaccine-mediated protection against Leishmania major. Nat Med. 2007; 13(7):843–850. doi:10.1038/nm1592.

10. Seder RA, Darrah PA, Roederer M. T-cell quality in memory and protection: Implications for vaccine design. Nat Rev Immunol. 2008; 8(4):247–258. doi:10.1038/nri2274.

11. Tarkowski A, Czerkinsky C, Nilsson LÅ, Nygren H, Ouchterlony Ö. Solid-phase enzyme-linked immunospot (ELISPOT) assay for enumeration of IgG rheumatoid factor-secreting cells. J Immunol Methods. 1984; 72(2):451–459. doi:10.1016/0022-1759(84)90013-9.

12. Newell EW, Sigal N, Bendall SC, Nolan GP, Davis MM. Cytometry by Time-of-Flight Shows Combinatorial Cytokine Expression and Virus-Specific Cell Niches within a Continuum of CD8+T Cell Phenotypes. Immunity. 2012; 36(1):142–152. doi:10.1016/j.immuni.2012.01.002.

13. Bendall SC, Simonds EF, Qiu P, et al. Single-Cell Mass Cytometry of Differential. Science (80-). 2011; 332(May):687–695. doi:10.1126/science.1198704.

14. Ullal A V., Peterson V, Agasti SS, et al. Cancer cell profiling by barcoding allows multiplexed protein analysis in fine-needle aspirates. Sci Transl Med. 2014; 6(219). doi :10.1126/scitranslmed.3007361.

15. Hughes AJ, Spelke DP, Xu Z, Kang C-C, Schaffer D V, Herr AE. Single-cell western blotting: Supplimentary. Nat Methods. 2014; 11(7):749–755. doi:10.1038/nmeth.2992.

16. Chattopadhyay PK, Gierahn TM, Roederer M, Love JC. Single-cell technologies for monitoring immune systems. Nat Immunol. 2014; 15(2):128–135. doi:10.1038/ni.2796.

17. Altelaar AFM, Munoz J, Heck AJR. Next-generation proteomics: Towards an integrative view of proteome dynamics. Nat Rev Genet. 2013; 14(1):35–48. doi:10.1038/nrg3356.

18. Ferguson A, Wang L, Altman RB, et al. Functional Dynamics within the Human Ribosome Regulate the Rate of Active Protein Synthesis. Mol Cell. 2015; 60(3):475–486. doi:10.1016/j.molcel.2015.09.013.

19. Tay S, Hughey JJ, Lee TK, Lipniacki T, Quake SR, Covert MW. Single-cell NF-B dynamics reveal digital activation and analogue information processing. Nature. 2010; 466(73 03):267–271. doi:10.1038/nature09145.

20. Han Q, Bagheri N, Bradshaw EM, Hafler DA, Lauffenburger DA, Love JC. Polyfunctional responses by human T cells result from sequential release of cytokines. Proc Natl Acad Sci. 2012; 109(5):1607–1612. doi:10.1073/pnas.1117194109.

21. Han Q, Bradshaw EM, Nilsson B, Hafler DA, Love JC. Multidimensional analysis of the frequencies and rates of cytokine secretion from single cells by quantitative microengraving. Lab Chip. 2010; 10(11):1391–1400. doi:10.1039/b926849a.

22. Kleppe M, Kwak M, Koppikar P, et al. JAK-STAT pathway activation in malignant and nonmalignant cells contributes to MPN pathogenesis and therapeutic response. Cancer Discov. 2015; 5(3):316–331. doi:10.1158/2159-8290.CD-14-0736.

23. Junkin M, Kaestli AJ, Cheng Z, et al. High-Content Quantification of Single-Cell Immune Dynamics. Cell Rep. 2016; 15(2):411–422. doi:10.1016/j.celrep.2016.03.033.

24. Lu Y, Chen JJ, Mu L, et al. High-Throughput Secretomic Analysis of Single Cells to Assess Functional Cellular Heterogeneity. Anal Chem. 2013; 85(4):2548–2556. doi:10.1021/ac400082e.

25. Lu Y, Xue Q, Eisele MR, et al. Highly multiplexed profiling of single-cell effector functions reveals deep functional heterogeneity in response to pathogenic ligands. Proc Natl Acad Sci. 2015; 112(7):E607–E615. doi:10.1073/pnas.1416756112.

26. Ma C, Fan R, Ahmad H, et al. A clinical microchip for evaluation of single immune cells reveals high functional heterogeneity in phenotypically similar T cells. Nat Med. 2011; 17(6):738–743. doi:10.1038/nm.2375.

27. Shi Q, Qin L, Wei W, et al. Single-cell proteomic chip for profiling intracellular signaling pathways in single tumor cells. Proc Natl Acad Sci. 2012; 109(2):419–424. doi:10.1073/pnas.1110865109.

28. Makamba H, Kim JH, Lim K, Park N, Hahn JH. Surface modification of poly(dimethylsiloxane) microchannels. Electrophoresis. 2003; 24(21):3607–3619. doi:10.1002/elps.200305627.

29. Gordon S, Taylor PR. Monocyte and macrophage heterogeneity. Nat Rev Immunol. 2005; 5(12):953–964. doi:10.1038/nri1733.

30. Mosser DM, Edwards JP. Exploring the full spectrum of macrophage activation. Nat Rev Immunol. 2008; 8(12):958–969. doi:10.1038/nri2448.

31. Balboa MA, Pérez R, Balsinde J. Amplification mechanisms of inflammation: paracrine stimulation of arachidonic acid mobilization by secreted phospholipase A2 is regulated by cytosolic phospholipase A2-derived hydroperoxyeicosatetraenoic acid. J Immunol. 2003; 171(2):989–994. doi:10.4049/JIMMUN0L.171.2.989.

32. Grkovich A, Johnson CA, Buczynski MW, Dennis EA. Lipopolysaccharide-induced cyclooxygenase-2 expression in human U937 macrophages is phosphatidic acid phosphohydrolase-1-dependent. J Biol Chem. 2006; 281(44):32978–32987. doi:10.1074/jbc.M605935200.

33. Xue Q, Lu Y, Eisele MR, et al. Analysis of single-cell cytokine secretion reveals a role for paracrine signaling in coordinating macrophage responses to TLR4 stimulation.

34. Garrelds IM, van Hal PT, Haakmat RC, Hoogsteden HC, Saxena PR, Zijlstra FJ. Time dependent production of cytokines and eicosanoids by human monocytic leukaemia U937 cells; effects of glucocorticosteroids. Mediators Inflamm. 1999; 8(4-5):229–235. doi:10.1080/09629359990397.

35. Amir EAD, Davis KL, Tadmor MD, et al. ViSNE enables visualization of high dimensional single-cell data and reveals phenotypic heterogeneity of leukemia. Nat Biotechnol. 2013; 31(6):545–552. doi:10.1038/nbt.2594.

36. Ma C, Fan R, Elitas M. Single Cell Functional Proteomics for Assessing Immune Response in Cancer Therapy: Technology, Methods, and Applications. Front Oncol. 2013; 3(May):1–7. doi:10.3389/fonc.2013.00133.

37. Larsen M, Sauce D, Arnaud L, Fastenackels S, Appay V, Gorochov G. Evaluating Cellular Polyfunctionality with a Novel Polyfunctionality Index. Hoshino Y, ed. PLoS One. 2012; 7(7):e42403. doi:10.1371/journal.pone.0042403.

38. Macosko EZ, Basu A, Satija R, et al. Highly parallel genome-wide expression profiling of individual cells using nanoliter droplets. Cell. 2015; 161(5):1202–1214. doi:10.1016/j.cell.2015.05.002.

39. Gaudillière B, Fragiadakis GK, Bruggner R V., et al. Clinical recovery from surgery correlates with single-cell immune signatures. Sci Transl Med. 2014; 6(255). doi:10.1126/scitranslmed.3009701.

40. Irish JM, Kotecha N, Nolan GP. Mapping normal and cancer cell signalling networks: towards single-cell proteomics. Nat Rev Cancer. 2006; 6(2):146–155. doi:10.1038/nrc1804.

41. Meacham CE, Morrison SJ. Tumour heterogeneity and cancer cell plasticity. Nature. 2013; 501(7467):328–337. doi:10.1038/nature12624.

42. Petricoin EF, Zoon KC, Kohn EC, Barrett JC, Liotta LA. Clinical proteomics: Translating benchside promise into bedside reality. Nat Rev Drug Discov. 2002; 1(9):683–695. doi:10.1038/nrd891.

43. Varadarajan N, Kwon DS, Law KM, et al. Rapid, efficient functional characterization and recovery of HIV-specific human CD8+ T cells using microengraving. Proc Natl Acad Sci. 2012; 109(10):3885–3890. doi:10.1073/pnas.1111205109.

44. Heath JR, Ribas A, Mischel PS. Single-cell analysis tools for drug discovery and development. Nat Rev Drug Discov. 2015; advance on(3):204–216. doi:10.1038/nrd.2015.16.

45. Macosko EZ, Basu A, Satija R, et al. Highly Parallel Genome-wide Expression Profiling of Individual Cells Using Nanoliter Droplets. Cell. 2015; 161(5): 1202–1214. doi:10.1016/j.cell.2015.05.002.

46. Satija R, Farrell JA, Gennert D, Schier AF, Regev A. Spatial reconstruction of single-cell gene expression data. Nat Biotechnol. 2015; 33(5):495–502. doi:10.1038/nbt.3192.

47. Luo W, Friedman MS, Shedden K, Hankenson KD, Woolf PJ. GAGE: generally applicable gene set enrichment for pathway analysis. BMC Bioinformatics. 2009; 10:161. doi:10.1186/1471-2105-10-161.

48. Luo W, Brouwer C. Pathview: an R/Bioconductor package for pathway-based data integration and visualization. Bioinformatics. 2013; 29(14):1830–1831. doi :10.1093/bioinformatics/btt285.

