## Supporting Information for "Multiplexed, sequential secretion analysis of the same single cells reveals distinct effector response dynamics dependent on the initial basal state"

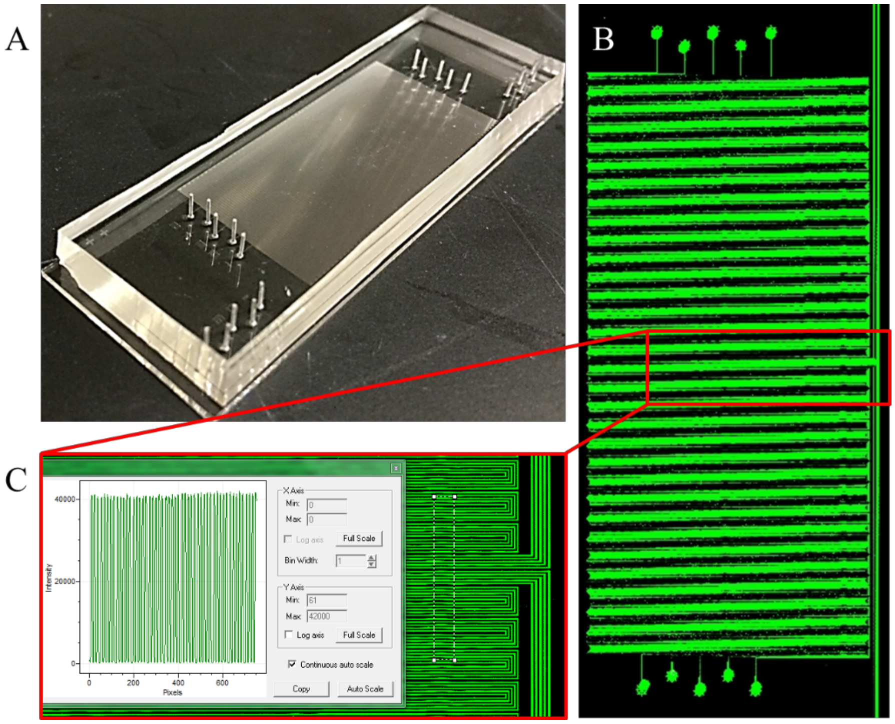


**Figure S1.** Evaluation of the protein coating uniformity flow-patterned with antibody barcode microchannel. A) Optical image showing the PDMS microchip for antibody flow patterning; B) whole glass slide fluorescence scanning; (C) magnified view of fluorescence intensity across the flow patterned poly-L-lysine slide revealed excellent uniformity of the immobilized proteins (FITC-BSA) in two separated paths, which ensures the validity of using this high-density barcode array technology to assess single cell heterogeneity.

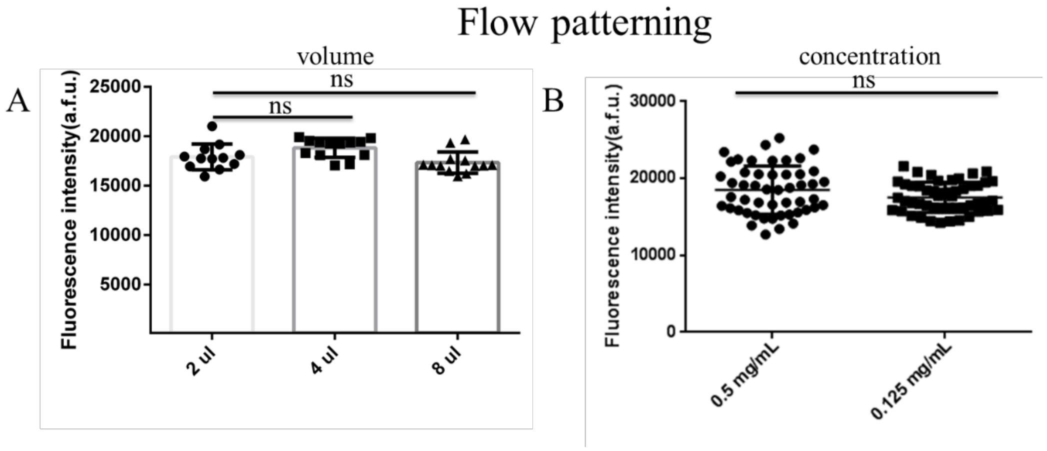


**Figure S2.** Evaluation of the antibody flow patterning conditions. A, different volume of antibodies; B, different antibody dilutions. The results showed no significant differences with varied flow patterning conditions.

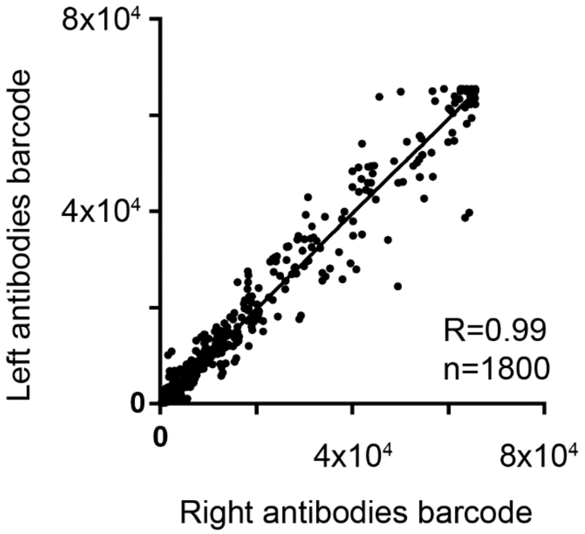


**Figure S3.** Characterization of the fluorescence detection results based on left and right antibodies barcode, which showed nice correlation between them.

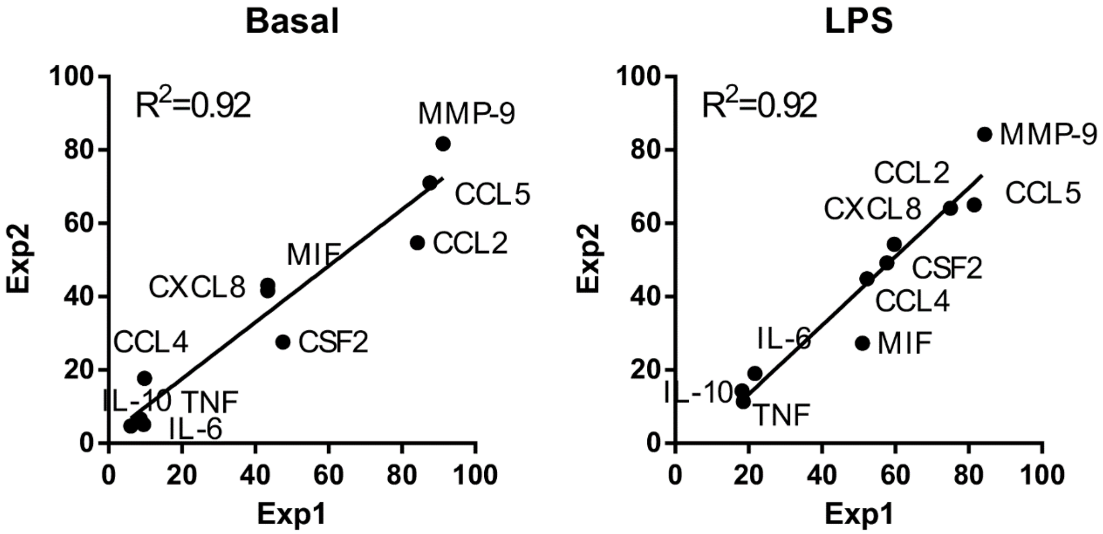


**Figure S4.** Validation of reproducibility of the single-cell barcode chip. Cytokine secretion frequencies (10 proteins) with U937 derived macrophages from two independent experiments (basal and LPS) showed high, linear correlation ($R^2$ = 0.92).

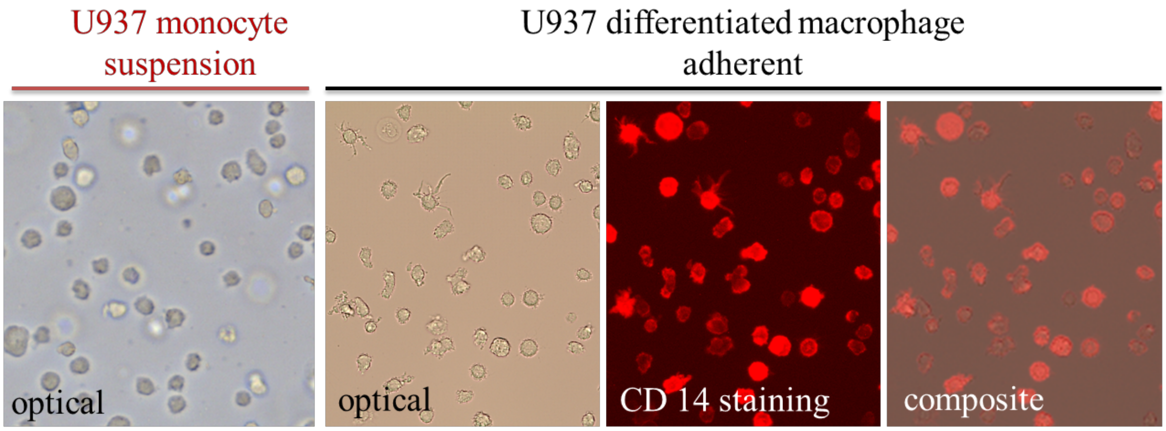


**Figure S5.** Images showing differentiation of U937 monocyte cells. Monocyte cells became adherent when differentiated into macrophage cells with 50 ng/mL PMA for 48 hours.

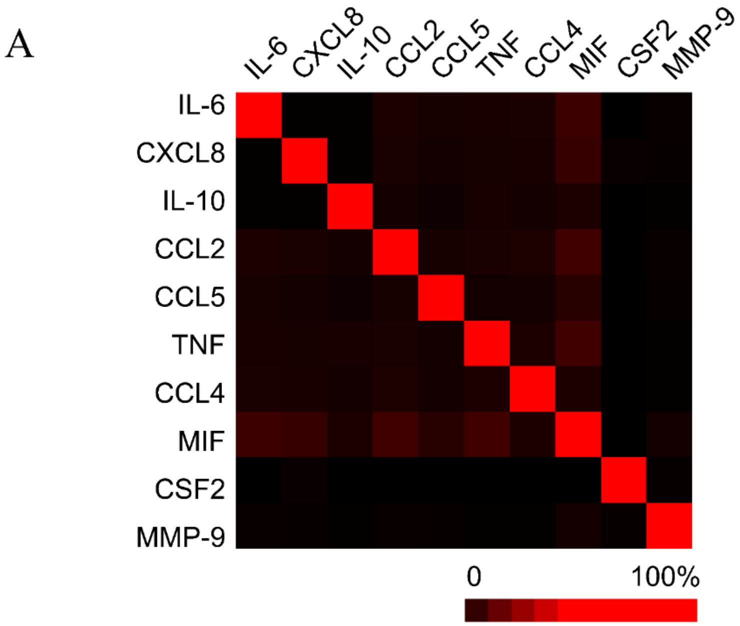


B

| Proteins | IL-6 | CXCL8 | IL-10 | CCL2 | CCL5 | TNF | CCL4 | MIF | CSF2 | MMP-9 |
|---|---|---|---|---|---|---|---|---|---|---|
| IL-6 | 100.0% | | | | | | | | | |
| CXCL8 | 1.1% | 100.0% | | | | | | | | |
| IL-10 | 0.9% | 0.9% | 100.0% | | | | | | | |
| CCL2 | 5.6% | 5.4% | 4.0% | 100.0% | | | | | | |
| CCL5 | 4.4% | 3.6% | 2.9% | 4.4% | 100.0% | | | | | |
| TNF | 4.7% | 4.5% | 4.9% | 5.2% | 3.7% | 100.0% | | | | |
| CCL4 | 5.2% | 4.7% | 4.0% | 5.8% | 4.2% | 5.7% | 100.0% | | | |
| MIF | 11.8% | 10.7% | 5.6% | 12.9% | 7.7% | 12.4% | 5.6% | 100.0% | | |
| CSF2 | 0.3% | 1.9% | 0.2% | 0.2% | 0.2% | 0.2% | 0.1% | 0.3% | 100.0% | |
| MMP-9 | 1.7% | 1.2% | 0.7% | 1.6% | 1.3% | 0.7% | 0.6% | 3.7% | 1.2% | 100.0% |

**Figure S6.** Cross-reactivity check for all proteins (A: heatmap; B: raw values). Most of the antibodies used in this study are monoclonal antibodies to ensure good specificity and reduce cross reactivity. The test was conducted by spiking a single antigen (recombinant protein) in a solution applied to a full capture antibody microarray containing all capture antibodies, followed by detection with a mixture of all detection Ab

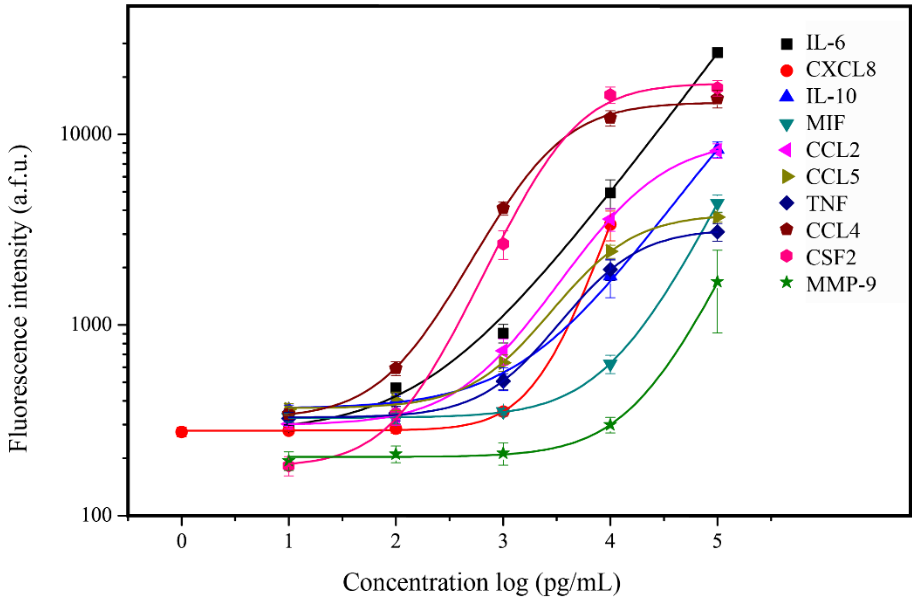


**Figure S7.** Calibration curves obtained with corresponding recombinant proteins (x, y axes: log-scaled, the intensities were averaged from 16 spots for each data).

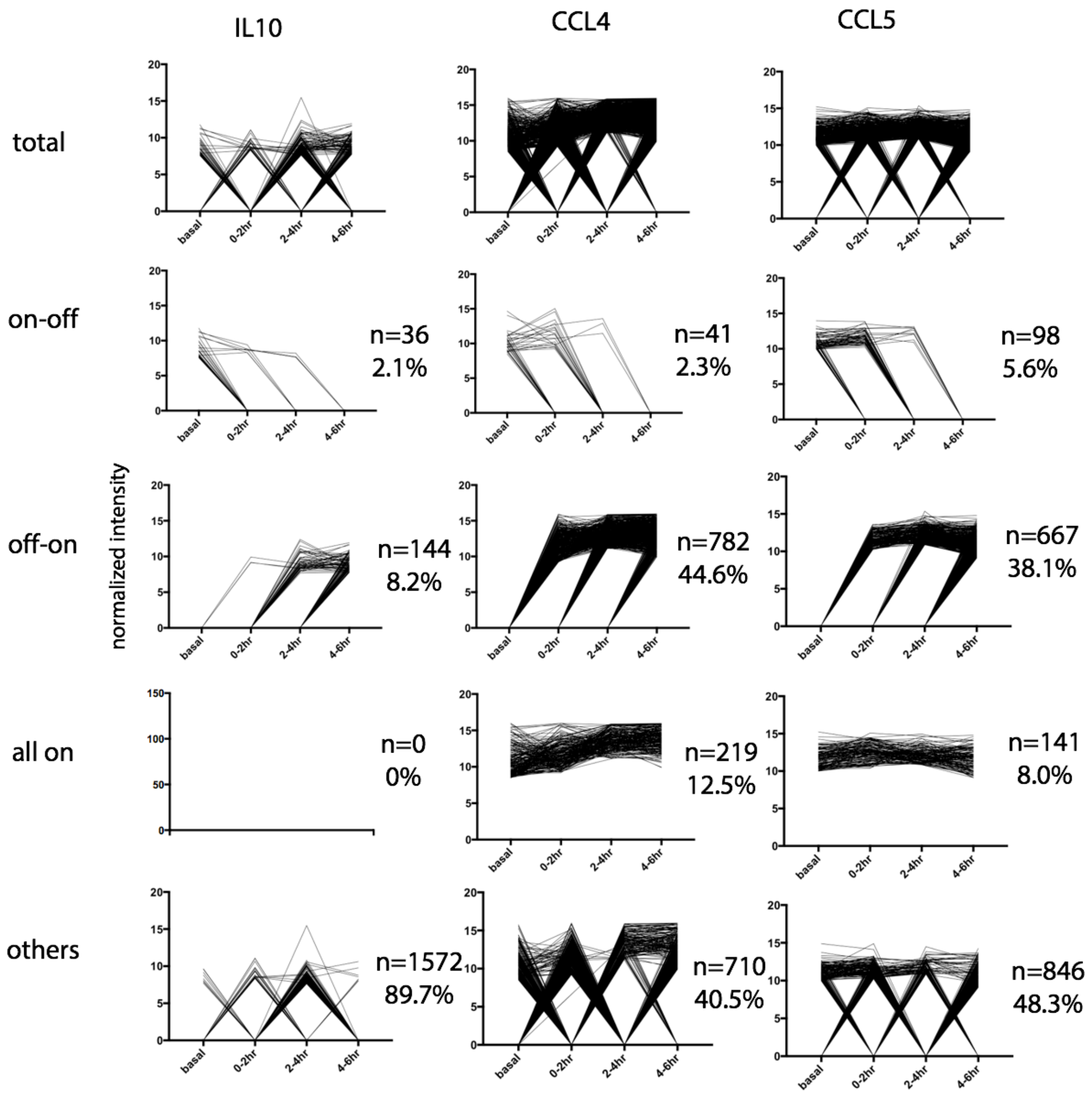


**Figure S8.** Classification of single cells based on their protein secretion dynamics.

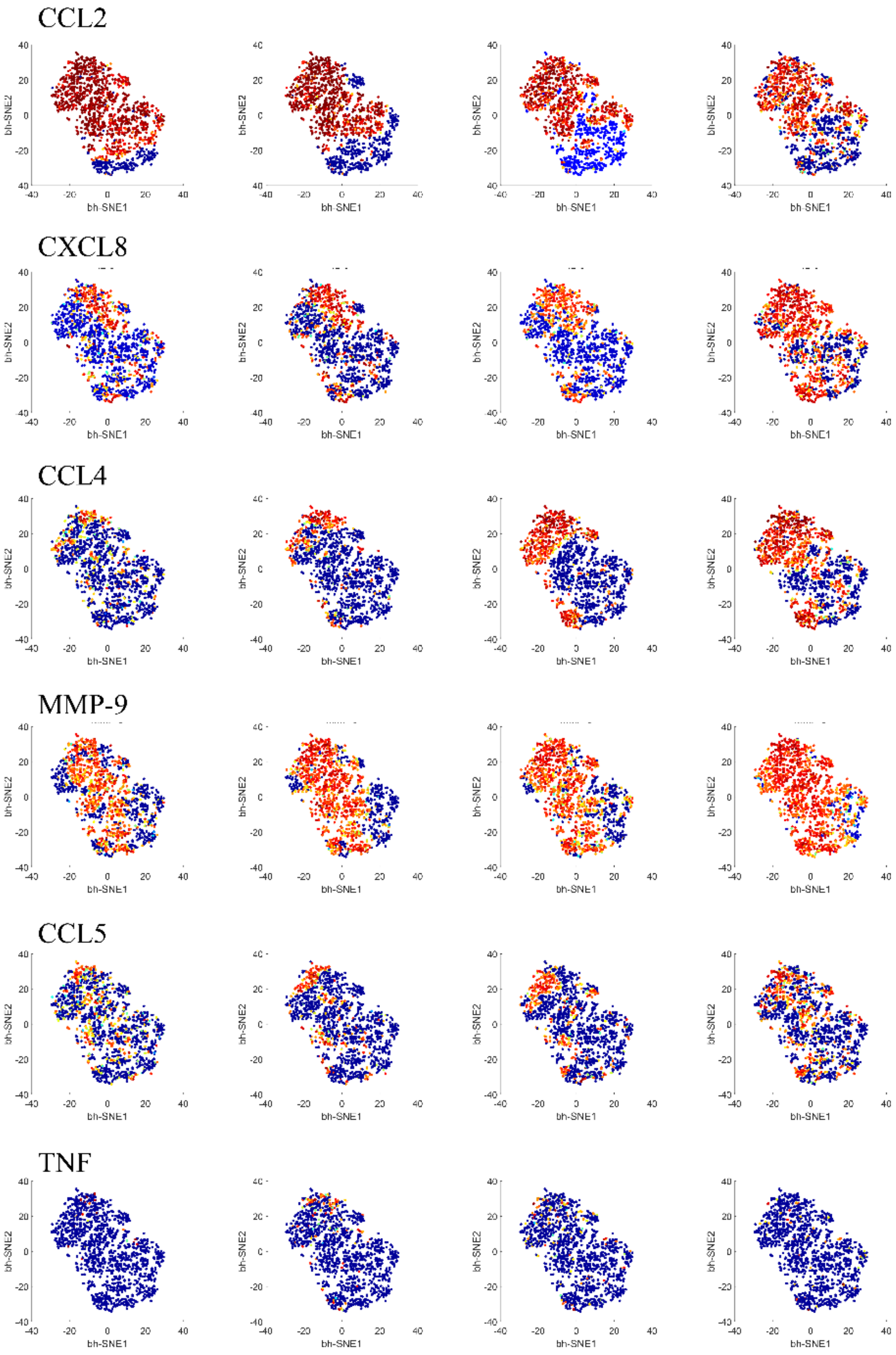
CCL2
CXCL8
CCL4
MMP-9
CCL5
TNF
bh-SNE1
bh-SNE2

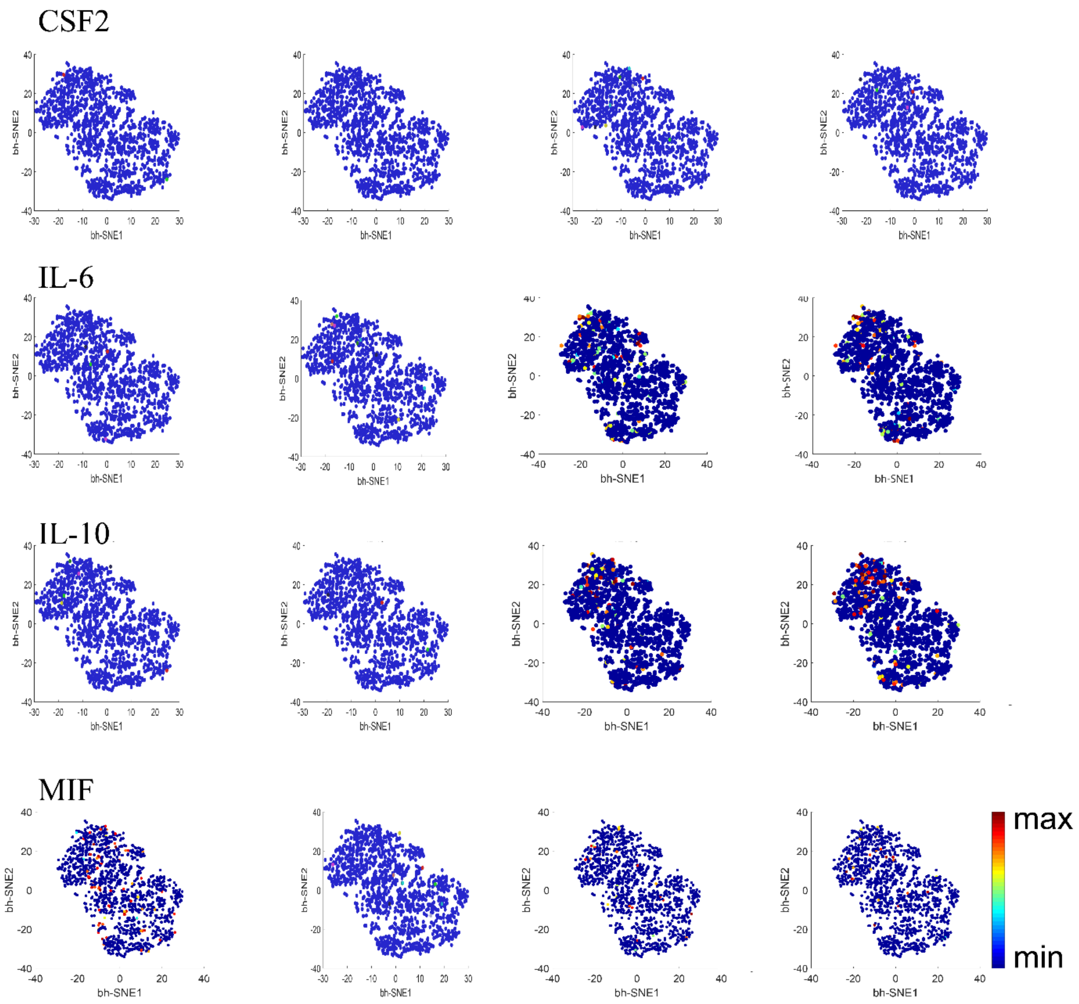


**Fig. S9** viSNE maps showing the distribution of each protein at different time points.

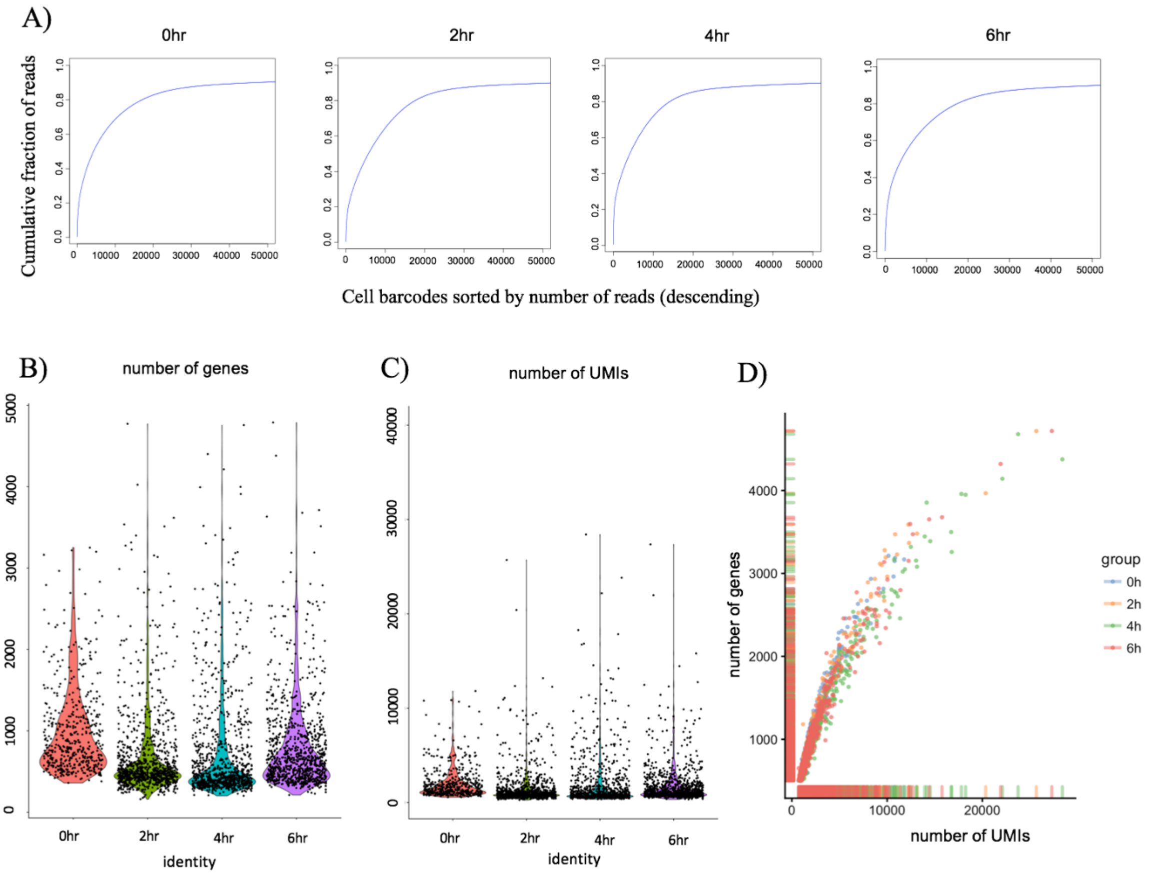


**Figure S10.** Quality control of single-cell RNA-Seq data. A) Saturation curve of sequencing reads. The deflection point of the curve shows the approximate number of cells sequenced. B) Violin plots showing distribution of number of genes detected from each single cell. C) Violin plots showing distribution of number of UMIs detected from each single cell. D) correlation between number of genes and UMIs detected from each single cell.

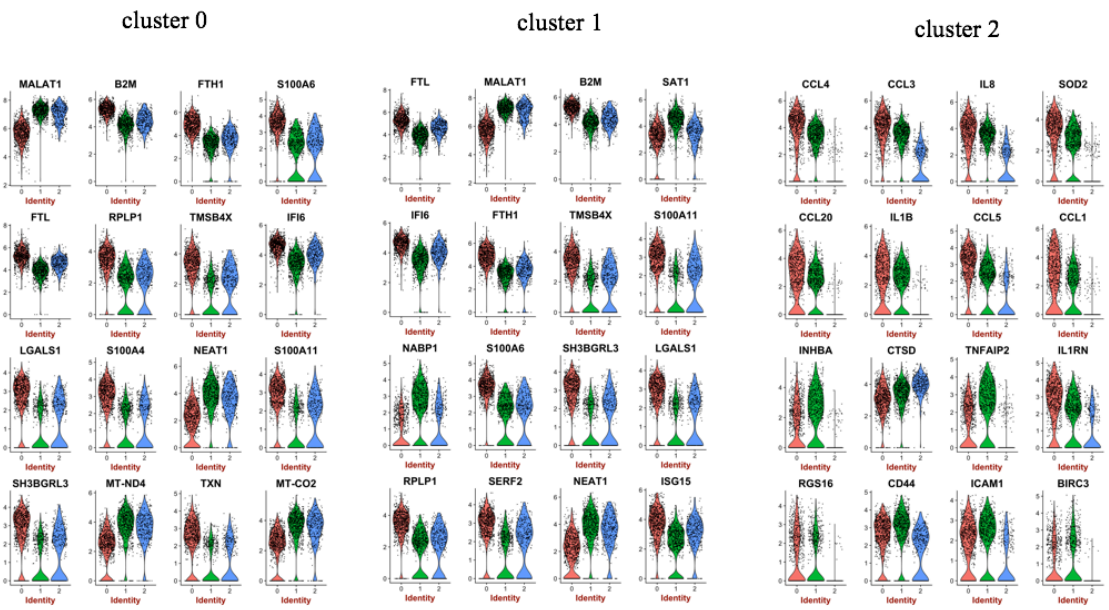


**Figure S11.** Top 16 marker genes of the three clusters, respectively.

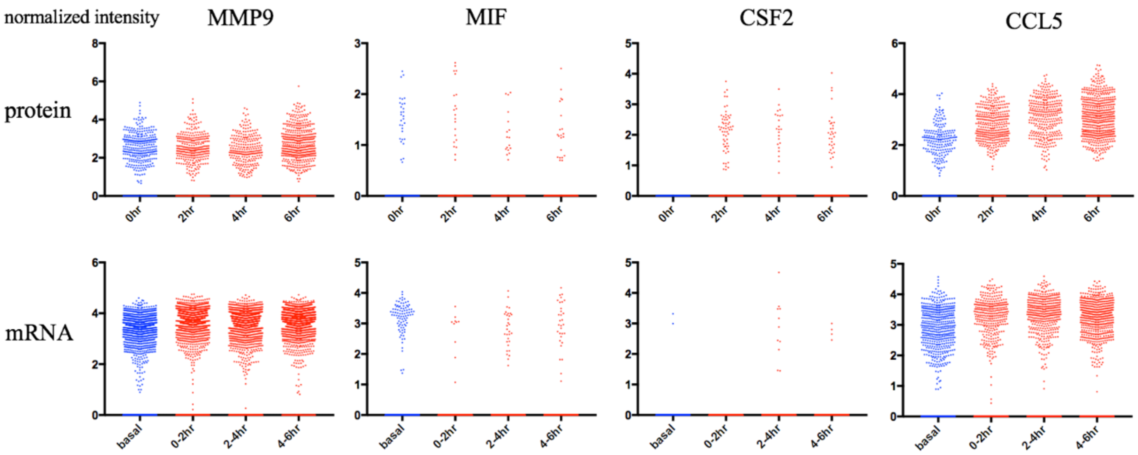


**Figure S12.** Comparative study of single-cell RNA expression and protein secretions.

**Supplementary Table S1:** Summary of antibodies (name, clone, company, catalog) used in this study (most are monoclonal antibodies).

| ***Protein*** | **Capture antibody Isotype/clone/vendor/catalog** | **Detection antibody Isotype/clone/vendor /catalog** |
|---|---|---|
| ***IL6*** | Rat IgG2a, κ /MQ2-39C3/Biolegend/501204 | Rat IgG1, κ /MQ2-13A5/Biolegend/ 501102 |
| ***CXCL8*** | Mouse IgG1, κ /H8A5/Biolegend/511502 | Mouse IgG1, κ /E8N1/Biolegend/ 511402 |
| ***IL10*** | Rat IgG2a, κ/JES3-12G8/Biolegend/ 501504 | Rat IgG1, κ /JES3-9D7/Biolegend/ 501402 |
| ***MIF*** | Mouse IgG1 Kappa/2A10-4D3/Abnova/H00004282-M01 | Mouse IgG1 Kappa/2A10-4D3/Abnova/H00004282-M01 |
| ***CCL2*** | Mouse IgG1, κ /5D3-F7/Biolegend/502607 | Armenian Hamster IgG /2H5/Biolegend/ 505902 |
| ***CCL5*** | Mouse IgG2b kappa/VL1/Invitrogen/AHC1052 | Mouse IgG1 /21418/RD/MAB678 |
| ***TNF*** | Mouse IgG1, κ /Mab11/Biolegend /502902 | Mouse IgG1, κ /MAb1/Biolegend/502802 |
| ***CCL4*** | Mouse IgG1 kappa/A174E 18A7/Invitrogen/AHC6114 | Mouse IgG2B /24006/RD/MAB271 |
| ***CSF2*** | Rat IgG2a, κ /BVD2-23B6/Biolegend/502202 | Rat IgG2a, κ /BVD2-21C11/Biolegend/502304 |
| ***MMP9*** | Mouse IgG1 /36020/RD/MAB936 | Goat IgG /polyclonal/RD/BAF911 |

**Supplementary Table S2:** Reagents used in single-cell RNA-Seq experiment.

| *Reagents* | Supplier | Catalog # |
|---|---|---|
| ***Barcoded microparticles*** | Chemgenes | N/A |
| ***Maxima H-Reverse Transcriptase*** | Thermo | EP0753 |
| ***dNTP mix*** | Clontech | 639125 |
| ***RNase inhibitor*** | Lucigen | 30281-2 |
| ***Perfluorooctanol*** | Sigma | 370533 |
| ***Exonuclease I*** | NEB | M0293L |
| ***KAPA Hifi HotStart Readymix*** | KAPA BioSystems | KK2602 |
| ***Nextera XT DNA sample preparation kit*** | Illumina | FC-131-1096 |
| ***Ampure XP beads*** | Backman Coulter | A63882 |
